# Mapping proteolytic neo-N termini at the surface of living cells

**DOI:** 10.1101/2020.02.23.961904

**Authors:** Amy M. Weeks, James R. Byrnes, Irene Lui, James A. Wells

## Abstract

N terminomics is a powerful strategy for profiling proteolytic neo-N termini, but its application to cell surface proteolysis has been limited by the low relative abundance of plasma membrane proteins. Here we apply plasma membrane-targeted subtiligase variants to efficiently and specifically capture cell surface N termini in live cells. Using this approach, we sequenced 807 cell surface N termini and quantified changes in their abundance in response to stimuli that induce proteolytic remodeling of the cell surface proteome. This technology will facilitate greater understanding of extracellular protease biology and reveal neo-N termini biomarkers and targets in disease.

## Main text

Proteolysis is a key post-translational modification that controls the function, localization, and degradation of nearly all proteins^1,2^. More than 500 proteases are encoded in the human genome, but many of their biological functions remain incompletely understood because few of their substrates are known. Several methods for selective isolation of protein N-terminal peptides, including those arising from proteolytic cleavage events, have been developed^3-6^, enabling global analysis of cellular proteolysis in response to biological perturbations. These strategies exploit the unique chemical structure and reactivity of the protein N terminus compared to other biological amines to isolate unblocked N termini by either positive enrichment or by depletion of internal peptides following protease digestion. The isolated N-terminal peptides can then be analyzed by tandem mass spectrometry (LC-MS/MS), enabling sequencing of protease cleavage sites with single amino acid resolution. While these techniques have proven powerful to identify protease substrates, they provide limited coverage of cell surface proteins, which often escape detection by mass spectrometry due to their low abundance relative to cytosolic and cytoskeletal proteins^7,8^. Cell surface proteolysis is crucial for cell-cell communication and is often dysregulated in disease. New techniques are therefore needed for identification of protease cleavage sites in plasma membrane proteins to facilitate biomarker and target discovery.

Subtiligase is a rationally designed variant of the serine protease subtilisin that harbors two key mutations (S221C, P225A) that enable it to catalyze a ligation reaction between a peptide ester and the N-terminal *α*-amine of a peptide or protein^9^. Based on this activity, subtiligase has been applied broadly as a tool for N-terminal modification to enable selective enrichment of N-terminal peptides and their identification and quantification by LC-MS/MS (**Fig. 1a**)^3,10^. This strategy, known as subtiligase N terminomics, has uncovered thousands of protease cleavage events that are stimulated in the contexts of apoptosis^3^, inflammation^11^, bacterial^12^ and viral infection^13^, and protein trafficking^14^. However, cell surface proteins are under-represented in N terminomics datasets due to their low abundances (**Fig. 1b, Supplementary Dataset 1**)^15^. We sought to develop a method to target cell surface N termini while avoiding the low specificity and material losses that would result from traditional approaches to isolate the plasma membrane. Our approach is to target subtiligase activity to the surface of living cells, where membranes, protein complexes, and spatial relationships remain intact.

**Figure 1.**
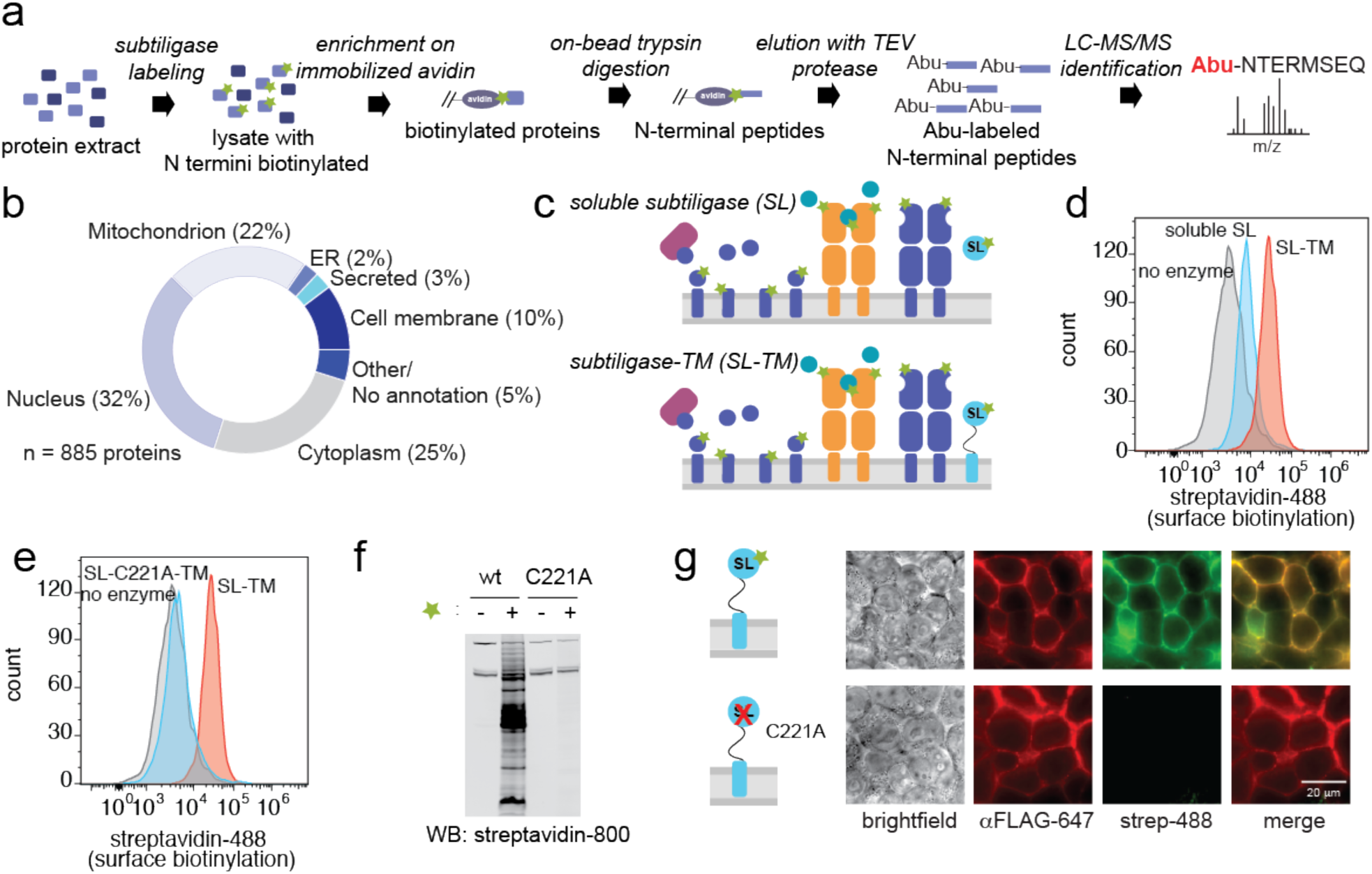
Restricting subtiligase activity to the cell surface. (a) Workflow for subtiligase N terminomics in cell lysate. Biotinylated peptide ester is represented by a green star. (b) Subcellular locations of N termini identified in a subtiligase N terminomics experiment performed using soluble subtiligase in cell lysate. (c) Approaches for labeling cell surface N termini. Top, addition of soluble subtiligase to live cells. Bottom, expression of subtiligase fused to the PDGFRβ chain (subtiligase-TM) in cells. (d) Streptavidin-488 flow cytometry demonstrates that subtiligase-TM (red) more efficiently biotinylates cell surface N termini compared to soluble subtiligase added to cells (cyan). No enzyme control is shown in grey. (e) Streptavidin-488 flow cytometry shows that robust cell surface biotinylation is observed with activate subtiligase-TM (red) and not the catalytically inactive C221A mutant (cyan). (f) A western blot of subtiligase-TM-expressing HEK293T cells shows that biotinylation activity is dependent on both active subtiligase and the presence of a biotinylated peptide ester substrate. No biotinylation is observed when the inactive subtiligase-C221A-TM mutant is used, or in the absence of biotinylated peptide ester substrate. (g) Fluorescence microscopy shows that subtiligase-TM expression (red) and biotinylation activity (green) are co-localized at the cell surface. No biotinylation activity is observed when the inactivate subtiligase-C221A-TM mutant is expressed.

We initially attempted to restrict subtiligase activity to the cell surface by adding the enzyme and a cell impermeable, negatively charged, biotinylated subtiligase substrate to a suspension of living cells (**Fig 1c**, top). Staining of the cells with streptavidin-Alexa Fluor 488 revealed modest biotinylation of the cell surface (**Fig. 1d**). We reasoned that the efficiency of cell surface N-terminal biotinylation could be increased by tethering subtiligase to the extracellular side of the plasma membrane (**Fig. 1c**, bottom). We therefore genetically targeted subtiligase to the desired location by fusing it to the transmembrane domain of the platelet-derived growth factor receptor beta chain (PDGFRβ-TM). We initiated labeling by adding the cell impermeable biotinylated substrate and measured cell surface biotinylation by flow cytometry. This experiment revealed a robust increase in biotinylation of approximately 10-fold compared to the soluble enzyme (**Fig. 1d**). In contrast, a catalytically inactive mutant of subtiligase (C221A) did not show any cell surface biotinylation activity (**Fig. 1e**). Streptavidin blot analysis of cell lysate demonstrated that many proteins were biotinylated in a manner that was dependent on both subtiligase activity and the presence of biotinylated subtiligase substrate (**Fig. 1f**), and that maximal biotinylation was achieved within 10 minutes (**Supplementary Fig. 1**). Fluorescence microscopy analysis of subtiligase expression and biotinylation activity showed that biotinylation and subtiligase activity were co-localized at the cell surface (**Fig. 1g**).

We next tested plasma membrane-targeted subtiligase in an MS N terminomics experiment (**Fig. 2a**). Proteins biotinylated by subtiligase were enriched on neutravidin beads, an on-bead trypsin digestion was performed to remove internal peptides, and N-terminal peptides were selectively eluted by cleavage of a TEV protease site incorporated into the subtiligase substrate. Following TEV cleavage, each subtiligase-modified N terminus retains an aminobutyric acid (Abu) mass tag for unequivocal identification. From four experiments, a total of 737 unique N termini from 371 unique proteins were identified (**Supplementary Dataset 3**). Subcellular location annotations for these proteins in Uniprot^16^ revealed that our N-terminal dataset was highly enriched for cell membrane and secreted proteins, with 249 of the identified proteins (62%) having a subcellular location annotation (cell membrane or secreted) suggesting that they would be present at the cell surface (**Fig. 2b, Supplementary Dataset 4**). To enable exploration of our datasets, we developed an <u>A</u>tlas of <u>S</u>ubtiligase-<u>C</u>aptured <u>E</u>xtracellular <u>N</u> <u>T</u>ermini (ASCENT, http://wellslab.org/ascent), for browsing and searching for subtiligase-captured cell surface N termini.

**Figure 2.**
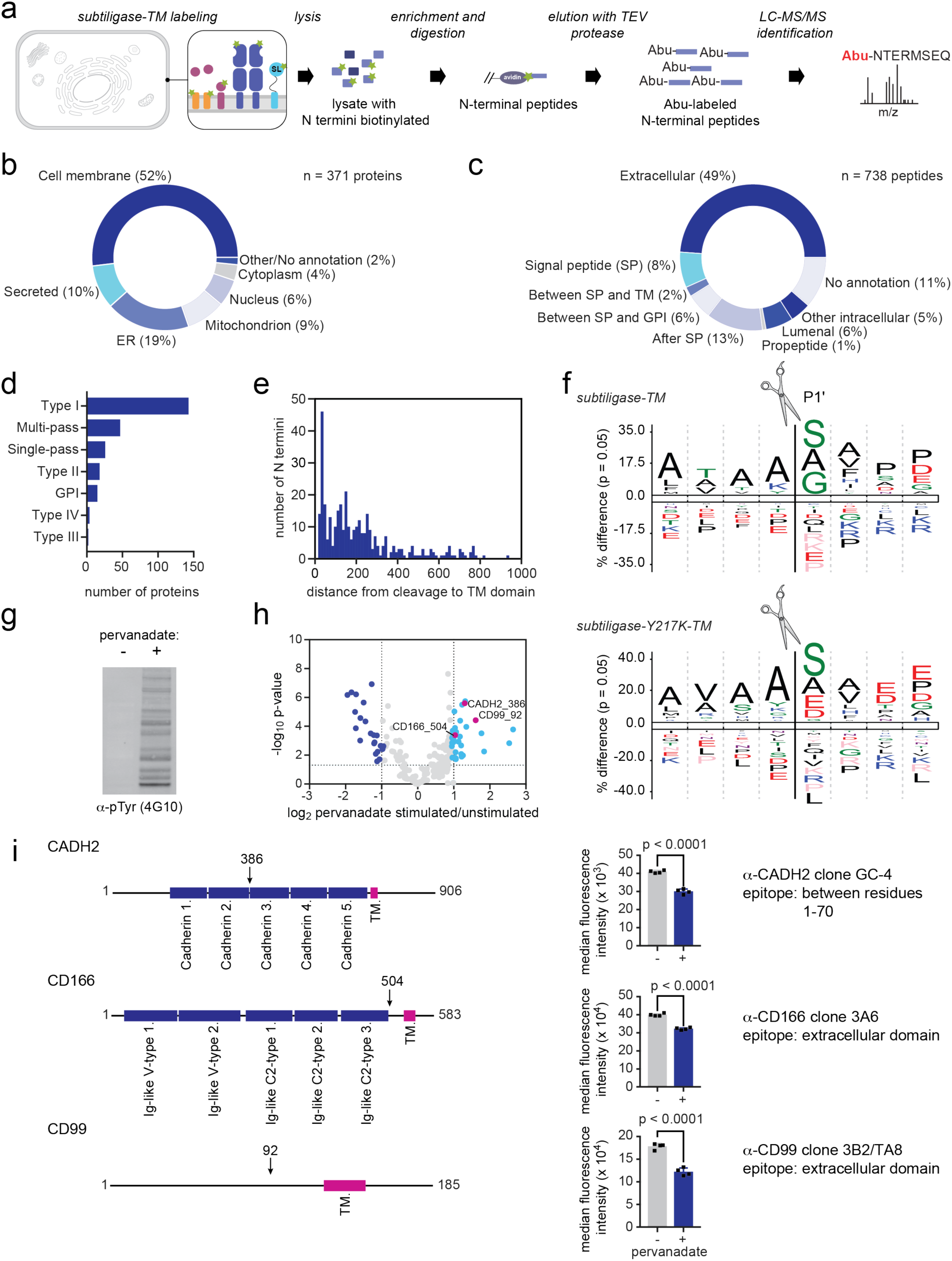
Cell surface N terminomics with subtiligase-TM. (a) Workflow for cell surface N terminomics with subtiligase-TM. (b) Subcellular locations of N termini identified in a subtiligase-TM N terminomics experiment performed in HEK293T cells.(c) Topological locations of N-terminal peptides identified in the subtiligase-TM N terminomics experiment. (d) Distribution of membrane protein types observed in the subtiligase-TM N terminomics experiment. (e) Distribution of distances between cleavage sites identified in the subtiligase N terminomics experiments and the corresponding transmembrane domains. (f) WebLogos for the N-terminal sequence specificity of wild-type subtiligase-TM and subtiligase-Y217K-TM. (g) Western blot showing an accumulation of phosphotyrosine upon pervanadate treatment of cells. (h) Volcano plot showing the log_2_-fold change of N termini measured by quantitative mass spectrometry in pervandate-treated versus untreated cells. The significance of the changes is indicated by -log_10_ p-value (y-axis). (i) Flow cytometry validation of changes in the N-terminal proteome observed in the mass spectrometry dataset. Monoclonal antibodies that bind epitopes N-terminal to the observed cleavage site were chosen such that a decrease in signal upon pervanadate treatment is expected.

Based on the design of the subtiligase-TM construct, we expect subtiligase activity to be restricted to the extracellular side of the plasma membrane and therefore to modify free N termini that are also located on the extracellular surface. To assess the specificity of subtiligase-TM for extracellular N termini, we mapped the N-terminal peptides captured with subtiligase-TM or with soluble subtiligase added to a lysate to topological domain annotations in Uniprot^16^ (**Fig. 2c, Supplementary Dataset 5**). Of the 737 N termini that we identified in the subtiligase-TM dataset, 360 (49%) mapped to annotated extracellular domains, while 23 (3%) mapped to cytoplasmic domains. For the remaining N termini that did not map to annotated extracellular domains, we mined Uniprot for other protein features that provided clues about their topological locations. The positions of many of the N termini relative to other protein features were consistent with an extracellular location. Among this group, 58 N termini corresponded to annotated signal peptide cleavages; 16 N termini were located between a signal peptide and a transmembrane domain; 42 N termini were located between a signal peptide and a GPI anchor; 6 N termini corresponded to annotated propeptide cleavage sites; and 92 N termini occurred between a signal peptide and the protein C terminus. The remaining N termini mapped to the ER lumen (45 sequences), other intracellular compartments (11 sequences), or had no annotation (84 sequences). In total, 79% of the identified N-terminal peptides were likely positioned on the extracellular surface during the live cell subtiligase-TM labeling experiment. In contrast, when we performed this analysis for N termini derived from the lysate dataset, of the 1693 N termini identified, 1% mapped to annotated extracellular domains and 4% of N termini have relationships to other features suggesting that they are derived from the extracellular surface (**Supplementary Fig. 2, Supplementary Dataset 6**). The majority of N termini in the lysate dataset (92%) have no topological annotation, suggesting that they are derived from proteins that are located entirely within a single compartment and are not associated with cellular membranes that would provide a topological orientation.

In order for subtiligase-TM to capture a neo-N terminus produced by cleavage of the extracellular domain of a protein, the neo-N terminus must remain associated with the cell surface. We therefore hypothesized that membrane protein types with their N-terminal domains on the extracellular surface (type I membrane proteins) would predominate over those with their C-terminal domains on the extracellular surface (type II membrane proteins). Consistent with this hypothesis, type I membrane proteins were overrepresented in our dataset compared to their abundance in the proteome (**Fig. 2d, Supplementary Dataset 7**). However, type II, type III, type IV, multipass, GPI-anchored, and other single-pass proteins were also observed in our dataset, suggesting that some proteolytically cleaved fragments that are not tethered to the plasma membrane may nonetheless stay associated with the cell.

We also hypothesized that the N-terminal peptides captured by subtiligase-TM would depend on the reach of subtiligase-TM from the membrane. Taking into account the length of the linker between subtiligase and the PDGFRβ-TM domain (estimated to be, on average, 23-27 Å^17^), as well as the size of subtiligase itself (estimated radius 20 Å^18^), we would expect that subtiligase should have the reach to modify even large extracellular domains up to ∼500 kDa (estimated radius 50 Å^18^). Consistent with this estimate, we observed subtiligase modification of extracellular N termini between 1 and 2585 amino acids away from their TM domains (**Fig. 2e**). The distribution of distances between observed N-terminal modification sites and TM domains was similar to the distribution of extracellular domain lengths across the proteome (**Supplementary Fig. 3, Supplementary Dataset 8**).

We next examined whether the use of subtiligase specificity mutants in the context of subtiligase-TM could increase coverage of N termini at the cell membrane in our experiment. We observed that the N-terminal specificity of subtiligase-TM (**Fig. 2f**) was similar to subtiligase added to cell lysate (**Supplementary Fig. 4**), with most of the N termini that were recovered having Ser, Gly, or Ala in the first position. However, when we introduced the Y217K mutation into subtiligase-TM, Glu and Asp N termini were also captured efficiently (**Fig. 2f**), while introduction of the Y217D mutation led to more efficient capture of His N termini (**Supplementary Fig. 5**). These results are consistent with our previous specificity data for these mutants^19^, and increase the total number of cell surface N termini that we identified to 807 (**Supplementary Dataset 9, Supplementary Dataset 10**).

We assessed the utility of subtiligase-TM for quantifying changes in levels of cell surface N termini by treating subtiligase-TM-expressing HEK293T cells with pervanadate, a covalent tyrosine phosphatase inhibitor that stimulates numerous cell surface proteolysis events^20^, or a vehicle control. Upon addition of pervanadate, we observed an accumulation of phosphotyrosine by Western blotting (**Fig. 2g**). We isotopically encoded the pervanadate-treated and untreated samples using SILAC, treated with a cell impermeable subtiligase substrate, and isolated biotinylated N-terminal peptides for identification and quantification by mass spectrometry. We identified 38 N termini that were significantly upregulated (>2-fold increase, p<0.05) upon pervanadate treatment, and 24 N termini that were significantly downregulated (>2-fold decrease, p<0.05) (**Fig. 2h**). Among the pervanadate-upregulated N termini were proteolytic products of proteins that are known sheddase targets, including cadherin-2 (CADH2, glypican-4 (GPC4), CD99, scavenger receptor class B member 1 (SCRB1/CD36), and CD166 (ALCAM)^21^. Notably, although these proteins were previously known to be shed from the cell surface, in many cases, the exact sites of proteolytic cleavage that our method identified were not previously known.

To validate our results, we used flow cytometry to examine loss of the domains that our mass spectrometry data indicated were cleaved. We selected three significantly upregulated N termini derived from proteins for which monoclonal antibodies with defined epitopes were available (CADH2_386, CD99_92, and CD166_504) (**Fig. 2i, Supplementary Fig. 6-11**). For the three proteins that we evaluated, flow cytometry confirmed the change that we observed in our mass spectrometry dataset.

Subtiligase-TM is a tool that combines the strengths of N terminomics methods such as terminal amine isotopic labeling of substrates (TAILS)^4^, combined fractional diagonal chromatography (COFRADIC)^5^, and traditional subtiligase N terminomics^3^, which provide information about the exact sites of proteolytic cleavage events. This complements methods such as secretome protein enrichment with click sugars (SPECS)^22^, which targets secreted or shed proteins but does not provide positional information nor identification of proteolytic events retained on the cell surface. These advantages make subtiligase-TM an attractive positive enrichment technique for identifying proteolytic cleavage sites in cell surface proteins at single amino acid resolution. Given the importance of extracellular proteolysis in health and disease we expect that this tool will be widely adopted by protease biologists and translational scientists to discover new biomarkers or neo-eptiopes for immunotherapy.

## Supporting information

Supplementary Results

## Acknowledgments

We thank S. Coyle, M. Ravalin, D. Sashital, and current and former members of the Wells laboratory for helpful discussions. This work was supported by NIH grant 5R01GM081051-09, the Chan Zuckerberg Biohub Investigatorship, and the Harry and Dianna Hind Professorship in Pharmaceutical Sciences (to J.A.W.). A.M.W. was supported by a Helen Hay Whitney Postdoctoral Fellowship (F-1112) and a Career Award at the Scientific Interface from the Burroughs Wellcome Fund (1017065).

## Author contributions

A.M.W. and J.A.W. designed the research. A.M.W., J.R.B., and I.L. performed the experiments. A.M.W. wrote the data analysis scripts and built the ASCENT database. A.M.W. and J.A.W. analyzed the data and interpreted results. A.M.W. and J.A.W. wrote the manuscript.

## Methods

### Cell lysate N terminomics

Cell lysate experiments were performed in Jurkat E6-1 cells. For each experiment, 250 million Jurkat cells were collected by centrifugation for 5 min at 300 × g and were washed twice with 50 mL PBS. Cells were resuspended in lysis buffer (400 mM tricine, pH 8, 4% (w/v) SDS, 100 μM PMSF, 100 μM AEBSF, 2.5 mM EDTA) and lysed by probe ultrasonication (20% amplitude, 10 cycles of 5s/1s on/off). Insoluble material was pelleted by centrifugation at 20,000g for 20 min at room temperature. The sample was boiled in the presence of 5 mM TCEP for 15 min to reduce disulfide bonds. Free cysteines were then alkylated in the presence of 10 mM iodoacetamide at room temperature in the dark for 1 h. Iodoacetamide was quenched by the addition of DTT to 25 mM final concentration. Triton X-100 was added to a final concentration of 2.5% (v/v) and the sample was diluted fourfold with water. Biotinylated subtiligase substrate 1 (**Supplementary Figure 9**) was added to a final concentration of 2.5 mM and the reaction was initiated by addition of stabiligase to a final concentration of 1 μM. The reaction mixture was incubated for 1 h at room temperature. After subtiligase labeling, biotinylated N-terminal peptides were enriched as described previously and analyzed by LC-MS/MS.

### Plasmid construction

Plasmids were constructed using Gibson cloning with *E. coli* XL10 as the cloning host. KOD Hot Start Polymerase (EMD Millipore) was used for PCR amplifications with the oligonucleotides listed in **Supplementary Table 9**. Plasmids were verified by Sanger sequencing (Quintara Biosciences). Subtiligase-TM was initially cloned into pCDNA3.2-Ig*k*-FLAG-GFP-PDGF TM (Wells lab) between the NdeI/BamHI sites. Subcloning was then performed as described below to construct vectors for lentiviral transduction.

#### pLX302-Igk-FLAG-Subtiligase-PDGF TM

A fragment encoding a fusion of the Ig*k* chain leader sequence, a Ca^2+^-independent variant of subtiligase, a 10x Gly-Ser linker, and the PDGF receptor β chain transmembrane domain was amplified from pCDNA3.2-Ig*k*-FLAG-GFP-PDGF TM using Ig*k*-FLAG-SL-PDGF F1 and R1 (**Supplementary Table 2**). The fragment was inserted between the BsrGI and NheI sites of pLX302 (Addgene #25896) using Gibson assembly.

#### Mutants of Igk-FLAG-Subtiligase-PDGF TM

Subtiligase mutants were constructed using overlap PCR with oligonucleotides listed in **Supplementary Table 2** and were inserted into pLX302 using Gibson assembly.

### Construction of cell lines expressing subtiligase-TM

To construct subtiligase-TM cell lines, HEK293T cells were lentivirally transduced. Lentivirus was produced by transfecting HEK293T cells at 80% confluence with a mixture of the transfer plasmid pLX302-Ig*k*-FLAG-Subtiligase-PDGF TM and the second-generation lentiviral packaging plasmids (pMD2.g and pCMV-R8.91) using FuGENE HD transfection reagent (Promega). After 6 h, the supernatant was removed and replaced with complete DMEM. After 72 h, the virus-containing supernatant was passed through a 0.45 μm PVDF filter and used directly for infection of HEK293T cells. HEK293T cells to be infected were grown to 80% confluence in a 6-well plate. Media was removed and replaced with a mixture of virus (1 mL), complete DMEM containing 8 μg/mL polybrene (1.5 mL), and complete DMEM (0.5 mL). After 24 h, the lentivirus-containing media was removed and replaced with complete DMEM. Puromycin was added to 2 μg/mL 72 h after transfection to select for transduced cells. Expression was validated by flow cytometry using Alexa Fluor 647-conjugated anti-DYKDDDDK tag antibody (BioLegend #637315).

### Flow cytometry analysis of subtiligase-TM activity

Cells were dissociated by incubation with Versene (0.04% EDTA in Ca^2+^/Mg^2+^-free PBS) and collected by centrifugation at 300 × g for 5 min. Cells were blocked with PBS containing 3% BSA and stained with Alexa Fluor 647-anti-DYKDDDDK (BioLegend) at 0.5 μg/mL or with Alexa Fluor 488-streptavidin (Life Technologies) at 1 μg/mL for 30 min at 4°C. Cells were washed three times with PBS containing 3% BSA, resuspended in PBS, and analyzed on a Beckman Coulter CytoFlex flow cytometer. Data were analyzed using FlowJo software.

### Immunofluorescence

HEK293T cells expressing subtiligase-TM and control cells were plated on glass-bottom, poly-D-lysine-coated imaging dishes. Dishes were incubated for 24 h at 37°C under 5% CO_2_ atmosphere. Cell were washed with PBS, fixed with 1% paraformaldehyde, and permeabilized with 0.1% Triton X-100. Cells were blocked with PBS containing 3% BSA, and Alexa Fluor 647-anti-DYKDDDDK (0.5 μg/mL) and Alexa Fluor 488-streptavidin (1 μg/mL) were added. After a 30 min incubation at room temperature, cells were washed three times with PBS containing 3% BSA, washed with PBS, and imaged on a Zeiss Axio Observer Z1 using a 63x oil objective. False-color images were produced using FIJI software.

### Subtiligase-TM N terminomics

For each experiment, one 500 cm^2^ dish of HEK293T cells expressing subtiligase-TM or a specificity variant was grown to ∼80% confluency. Media was removed and cells were washed once with 50 mL PBS. Versene (25 mL) was added and plates were incubated at 37°C for 10 min to allow cell dissociation to occur. Cells were harvested by centrifugation at 300 × g for 5 min, washed with 50 mL PBS, and transferred into a 1.5 mL microcentrifuge tube. Cells were washed three times with 1 mL of labeling buffer (100 mM tricine, pH 8, 150 mM NaCl) and then resuspended in labeling buffer containing 2.5 mM Tev Ester 6 (**Supplementary Figure 9**). Subtiligase labeling was allowed to proceed for 1 h at 4°C on a rotating mixer. Following labeling, cells were pelleted by centrifugation at 300 × g for 5 min and washed three times with PBS. Cells were resuspended in RIPA lysis buffer (50 mM Tris, pH 7.4, 150 mM NaCl, 1% NP-40, 0.5% sodium deoxycholate, 0.1% SDS) supplemented with Halt Protease and Phosphatase Inhibitor Cocktail (ThermoFisher Scientific). Lysis was completed by probe ultrasonication (20% amplitude, 10 cycles of 5 s on / 1 s off). Insoluble material was pelleted by centrifugation at 20,000 × g for 20 min at 4°C. Biotinylated proteins were enriched from the supernatant on High-Capacity Neutravidin Agarose resin (ThermoFisher Scientific) (0.5 mL of 50% resin slurry). The resin was washed with each of the following buffers: RIPA (10 × 800 μL), PBS with 1 M NaCl (10 × 800 μL), 100 mM ammonium bicarbonate (10 × 800 μL), 100 mM ammonium bicarbonate with 2 M urea (10 × 800 μL). Resin was resuspended in 1 mL 100 mM ammonium bicarbonate with 2 M urea and transferred to a 1.5 mL microcentrifuge tube. 1 M TCEP was added to a final concentration of 5 mM and the sample was incubated at room temperature on a rotating mixer for 30 min. 500 mM iodoacetamide was added to a final concentration of 10 mM and the resin was incubated for 1 h at room temperature in the dark. Resin was pelleted at 500 × g, washed with 100 mM ammonium bicarbonate with 2 M urea (3 × 800 μL), and resuspended in 1 mL 100 mM ammonium bicarbonate with 2 M urea. Sequencing grade modified trypsin (20 μg, Promega) was added and digestion was allowed to proceed at room temperature on a rotating mixer overnight. The resin was then washed with with each of the following buffers: 100 mM ammonium bicarbonate with 2 M urea (10 × 800 μL), 100 mM ammonium bicarbonate (10 × 800 μL), 4 M guanidinium hydrochloride (10 × 800 μL), 100 mM ammonium bicarbonate (10 × 800 μL), TEV elution buffer (100 mM ammonium bicarbonate, 2 mM DTT) (10 × 800 μL). The resin was resuspended in 0.5 mL TEV elution buffer and incubated with TEV protease (10 μg) overnight at room temperature to elute biotinylated peptides. The eluted peptides were collected by spinning through a spin filter to remove the resin. The eluate was adjusted to 5% TFA, incubated on ice for 10 min, and centrifuged at 20,000 × g at 4°C to pellet precipitated TEV protease. Peptides were desalted on C18 spin tips (ThermoFisher Scientific), dried by vacuum centrifugation, dissolved in 10 μL 0.1% formic acid/2% acetonitrile, and analyzed by LC-MS/MS.

### LC-MS/MS analysis

Peptides were injected onto an Acclaim PepMap RSLC column (75 μm x 15 cm, 2 μm particle size, 100 Å pore size, ThermoFisher Scientific) and analyzed using a Thermo Dionex UltiMate 3000 RSLCnano liquid chromatography system and a Thermo Q-Exactive Plus hybrid quadrupole-Orbitrap mass spectrometer. Samples were loaded onto the column over 15 min at 0.5 μL/min in mobile phase A (0.1% formic acid). Peptides were eluted at 0.3 μL/min using a linear gradient from mobile phase A to 40% mobile phase B (0.1% formic acid, 80% acetonitrile) over 125 min. Data-dependent acquisition was performed scanning a mass range from 300-1,500 m/z using Thermo Xcalibur software.

### Mass spectrometry data analysis

Thermo RAW files were converted to peaklists using MSConvert (Proteowizard). Peaklists were searched again the human SwissProt database using Protein Prospector (UCSF) with a false discovery rate of <1%. The parent ion tolerance was set at 6 ppm and the fragment ion tolerance was set at 20 ppm. Tryptic specificity was required only at the C terminus of peptides to enable identification of endogenous proteolytic cleavages, and two missed tryptic cleavages were allowed. Carbamidomethylation at Cys was set as a constant modification and aminobutyric acid (Abu) at peptide N termini, acetylation at protein N termini, oxidation at Met, pyroglutamate formation at N-terminal Gln, and Met excision at protein N termini were set as variable modifications.

### Subcellular location, topological, and membrane protein type annotations

Subcellular location annotations and Gene Ontology (GO) Cellular Component annotations were used to determine the subcellular locations of proteins identified by LC-MS/MS. Annotations were retrieved from the Uniprot flat file using custom Python scripts. Uniprot subcellular location annotations were used when available; if a Uniprot subcellular location annotation was not available, GO Cellular Component annotations were used.

### Analysis of distance between TM domains and cleavage sites

Type I membrane proteins were analyzed to determine the distance between the cleavage sites observed in the subtiligase-TM dataset and the transmembrane domain annotated in Uniprot using a custom Python script.

### Sequence logos for subtiligase, subtiligase-TM, and specificity variants

N termini captured with subtiligase, subtiligase-TM, or subtiligase-TM specificity variants, as well as bioinformatically inferred sequences C-terminal to the cleavage site, were analyzed for enrichment or de-enrichment of each amino acid using the standalone version of iceLogo software. The human SwissProt database was used as a reference set with random sampling.

### Pervanadate treatment of subtiligase-TM-expressing HEK293T cells

Pervandate treatment and SILAC analysis was performed for subtiligase-TM, subtiligase-Y217K-TM, and subtiligase-Y217D-TM. For each replicate of the SILAC experiment, one 500 cm^2^ dish of subtiligase-TM-expressing cells was grown in heavy SILAC DMEM and one dish was grown in light SILAC DMEM. Two replicates were performed in which the light cells were treat with pervandate and two replicates were performed in which the heavy cells were treated with pervanadate. Before each experiment, a 50 mM stock solution of pervanadate was prepared fresh by mixing equal volumes of 100 mM sodium orthovanadate (New England Biolabs) and 100 mM hydrogen peroxide. Media was removed from the cells and replaced with serum-free heavy or light DMEM. The pervanadate stock solution was diluted 1:1000 into the serum-free media for pervanadate treatment. For control cells, and equal volume of water was added. Treated and untreated cells were incubated for 2 h at 37°C under 5% CO_2_ atmosphere. After 2 h, media was removed, cells were washed with PBS, and versene was added to lift the cells. Cells were harvested by centrifugation at 300 × g for 5 min, resuspended in PBS, and counted. Equal numbers of light pervanadate-treated and heavy untreated, or heavy pervanadate-treated and light untreated cells were mixed together and labeled according to the subtiligase-TM method described above.

### SILAC data analysis

SILAC quantification was performed using Skyline software. The results of a Protein Prospector search were exported as a pepXML file. Because of inconsistencies in how Protein Prospector and Skyline assign the N-terminal Abu modification, the pepXML file was modified to be Skyline-compatible using a custom Python script. An mzXML-format peaklist was generated from the Thermo RAW files corresponding to each experiment using MSConvert (Proteowizard). The pepXML, mzXML, and RAW files were imported into Skyline, along with a FASTA format list of the identified peptides. The total MS1 area and the isotope dot product for each peptide were calculated using Skyline and a report was exported for further analysis. The mean experiment-to-control ratio for each N-terminal peptide was calculated using a custom Python script. An isotope dot product of >0.95 for both the heavy and light MS1 peaks was required for an identified peptide to be quantified. A t-test was performed to evaluate the significance of the observed changes in experiment-to-control ratio across the four replicate experiments, treating each experiment as an independent sample.

### Flow cytometry for SILAC data validation

HEK293T cells were treated with pervanadate or vehicle as described above. Cells were dissociated by incubation with Versene (0.04% EDTA in Ca^2+^/Mg^2+^-free PBS) and collected by centrifugation at 300 × g for 5 min. Cells were blocked with PBS containing 3% BSA and stained with either APC-conjugate anti-human CD99 antibody clone 3B2/TA8 (1:20 dilution, BioLegend 341307), PE anti-human CD166 clone 3A6 (1:20 dilution, BioLegend 343903), or mouse monoclonal anti-N-Cadherin clone GC-4 followed by AlexaFluor 647-conjugated goat anti-mouse antibody (1:200 dilution, ThermoFisher Scientific). Staining was performed for 30 min at 4°C. Cells were washed three times with PBS containing 3% BSA, resuspended in PBS, and analyzed on a Beckman Coulter CytoFlex flow cytometer. Data were analyzed using FlowJo software.

### Statistical analysis

Statistical tests for flow cytometry were performed using Prism 8 (Graphpad). To compare pervanadate-treated versus untreated cells, the median fluorescence intensity (MFI) for the relevant channel was calculated using FlowJo. P values were calculated using a two-tailed, unpaired t test. Error bars indicate the mean MFI ± s.d. for four cell culture replicates. Statistical tests for SILAC mass spectrometry data were performed using a custom Python script (available at https://doi.org/10.5281/zenodo.3678926). P values were calculated using scipy.stats.ttest_ind, a built-in function of the SciPy library. A two-sided test for the null hypothesis that the heavy and light samples have identical values was performed. Unless otherwise indicated, four cell culture replicates for each experiment were performed, with the light cells being pervanadate-treated for two of the replicates and the heavy cells being pervanadate-treated for two of the replicates. Fluorescence microscopy images are representative of experiments performed for three cell culture replicates.

### Data availability

All data generated or analyzed for this study are available within the paper and its associated supplementary information files, from the ASCENT database (http://wellslab.org/ascent), or from the corresponding author upon reasonable request. In addition, raw mass spectrometry data and search results have been deposited in the ProteomeXchange repository under the accession numbers listed in **Supplementary Table 1**.

### Code availability

Python scripts used for data analysis are available in a GitHub repository (https://doi.org/10.5281/zenodo.3678926) under the GNU General Public License v3.0.

