## Supplementary Results for "Mapping proteolytic neo-N termini at the surface of living cells"

**Supplementary Table 1.** List of supplementary mass spectrometry dataset information and ProteomeXchange Accession Numbers.

| *dataset* | *description* | *experiments* | *ProteomeXchange Accession #* |
|---|---|---|---|
| 2 | Subtiligase lysate N terminomics data | 2 | PXD017687 |
| 3 | Subtiligase-TM N terminomics data | 4 | PXD017664 |
| 9 | Subtiligase-Y217K-TM N terminomics data | 1 | PXD017667 |
| 10 | Subtiligase-Y217D-TM N terminomics data | 1 | PXD017668 |
| 11a | Subtiligase-TM pervanadate SILAC data | 4 | PXD017669 |
| 11b | Subtiligase-Y217K-TM pervanadate SILAC data | 4 | PXD017685 |
| 11c | Subtiligase-Y217D-TM pervanadate SILAC data | 3 | PXD017686 |

**Supplementary Table 2.** List of oligonucleotides used for plasmid construction and site-directed mutagenesis.

| *name* | *sequence* |
|---|---|
| IgK-FLAG-SL-PDGF F1 | ggctaactgtcgggatcaacaagtttGTACAAAAAAGTTGGCACCATGGAGACAGACACACTCCTGCTATGGG |
| IgK-FLAG-SL-PDGF R1 | tcgacttaacgcgccaccggttagcgctagctcattactaTCACCTGGGCTTCTTCTGCCACAG |
| SL M222A F1 | acagcggtacgtgcgcggcatctgcgcacg |
| SL M222A R1 | cgtgcgcagatgccgcgcacgtaccgctgt |
| SL M222A Y217K F1 | ccggtacggggcgaagagcggtacgtgcg |
| SL M222A Y217K R1 | cgcacgtaccgctcttcgccccgtaccgg |
| SL M222A Y217D F1 | cggtacggggcggacagcggtacgt |
| SL M222A Y217D R1 | acgtaccgctgtccgccccgtaccg |

**Supplementary Figure 1.** Timecourse of biotinylation by subtiligase-TM expressed on the surface of HEK293T cells. Cells were lysed at various timepoints and the extent of biotinylation was measured by Western blotting with streptavidin for subtiligase-TM (dark blue) and subtiligase-C221-TM (light blue).

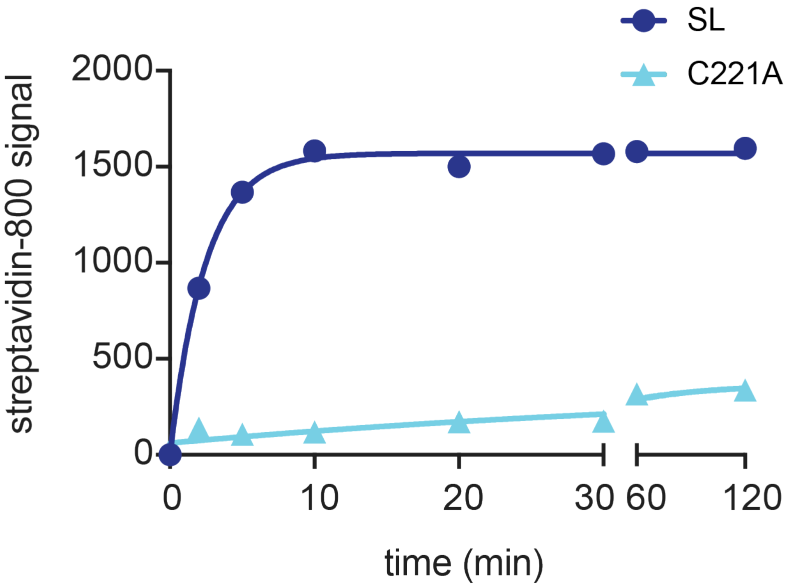

**Supplementary Figure 2.** Topological annotations for N-terminal peptides captured from Jurkat cell lysate using soluble subtiligase. Data for generating the donut plot can be found in Supplementary Dataset 6.

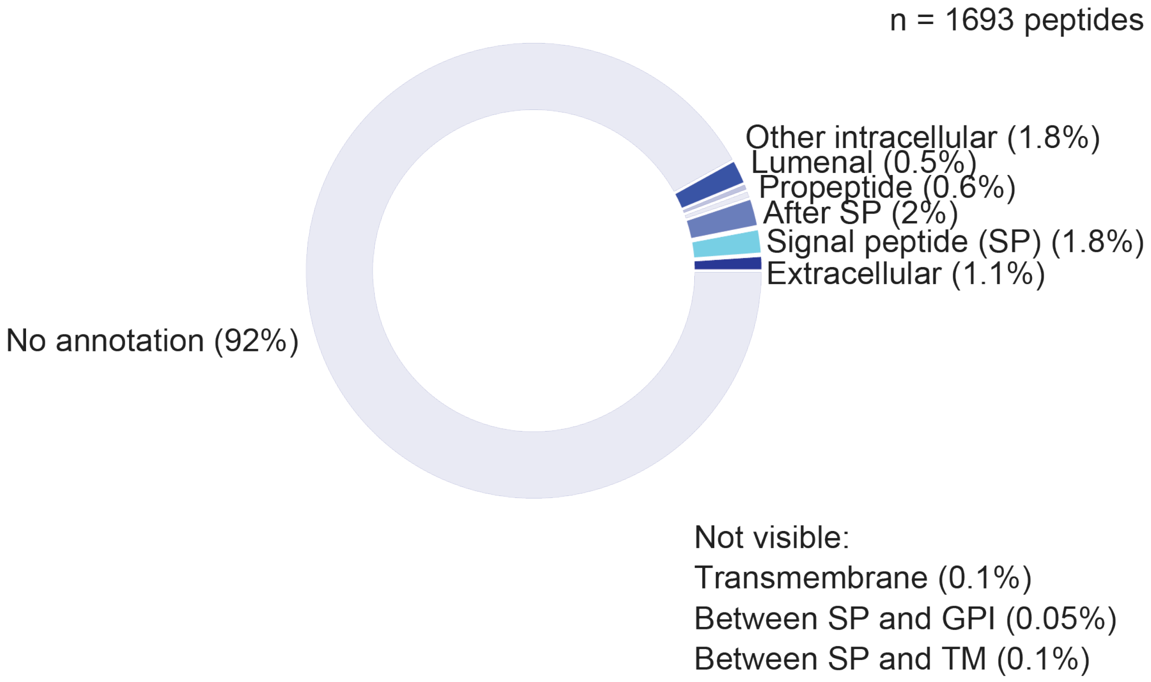

**Supplementary Figure 3.** Distribution of extracellular domain lengths across the human SwissProt database. The histogram below shows the frequency of extracellular domains of 0-1000 amino acids. Less than 10% of extracellular domains were longer than 1000 amino acids are not show on the histogram.

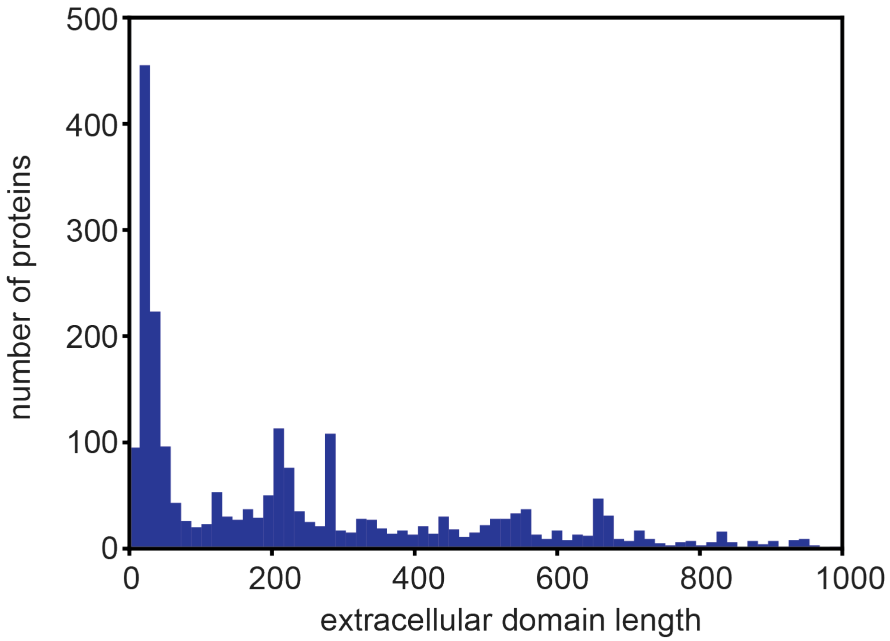

**Supplementary Figure 4.** IceLogo for N-terminal peptides captured using soluble subtiligase in Jurkat cell lysate. IceLogo was generated using stand-alone IceLogo software with random sampling of the human proteome as a reference.

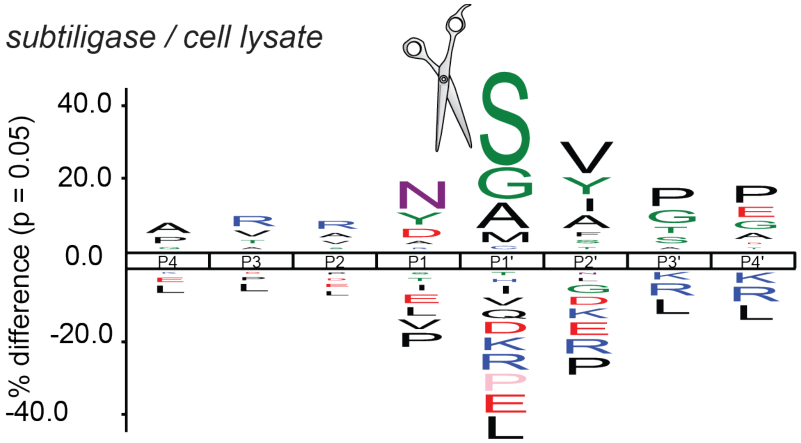

**Supplementary Figure 5.** IceLogo for N-terminal peptides captured using subtiligase-Y217D-TM expressed on the surface of HEK293T cells. IceLogo was generated using stand-alone IceLogo software with random sampling of the human proteome as a reference.

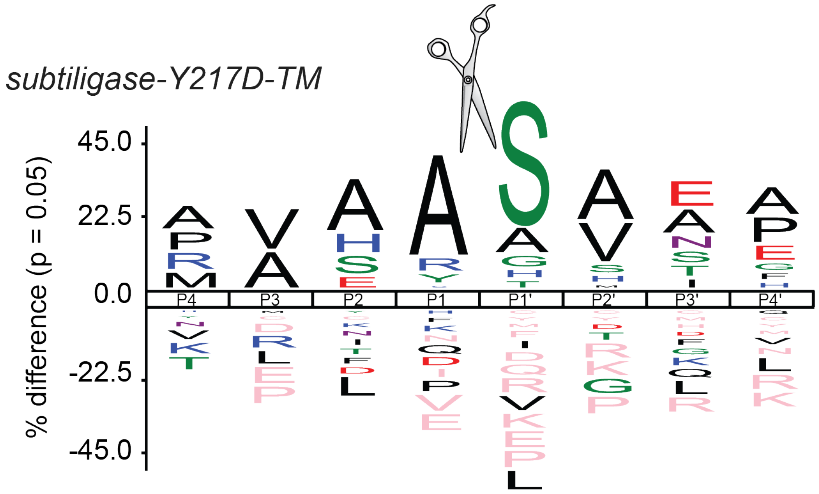

**Supplementary Figure 6.** Flow cytometry gating used for analysis of CADH2 levels on the surface of HEK293T cells in the (a) absence or (b) presence of pervanadate.

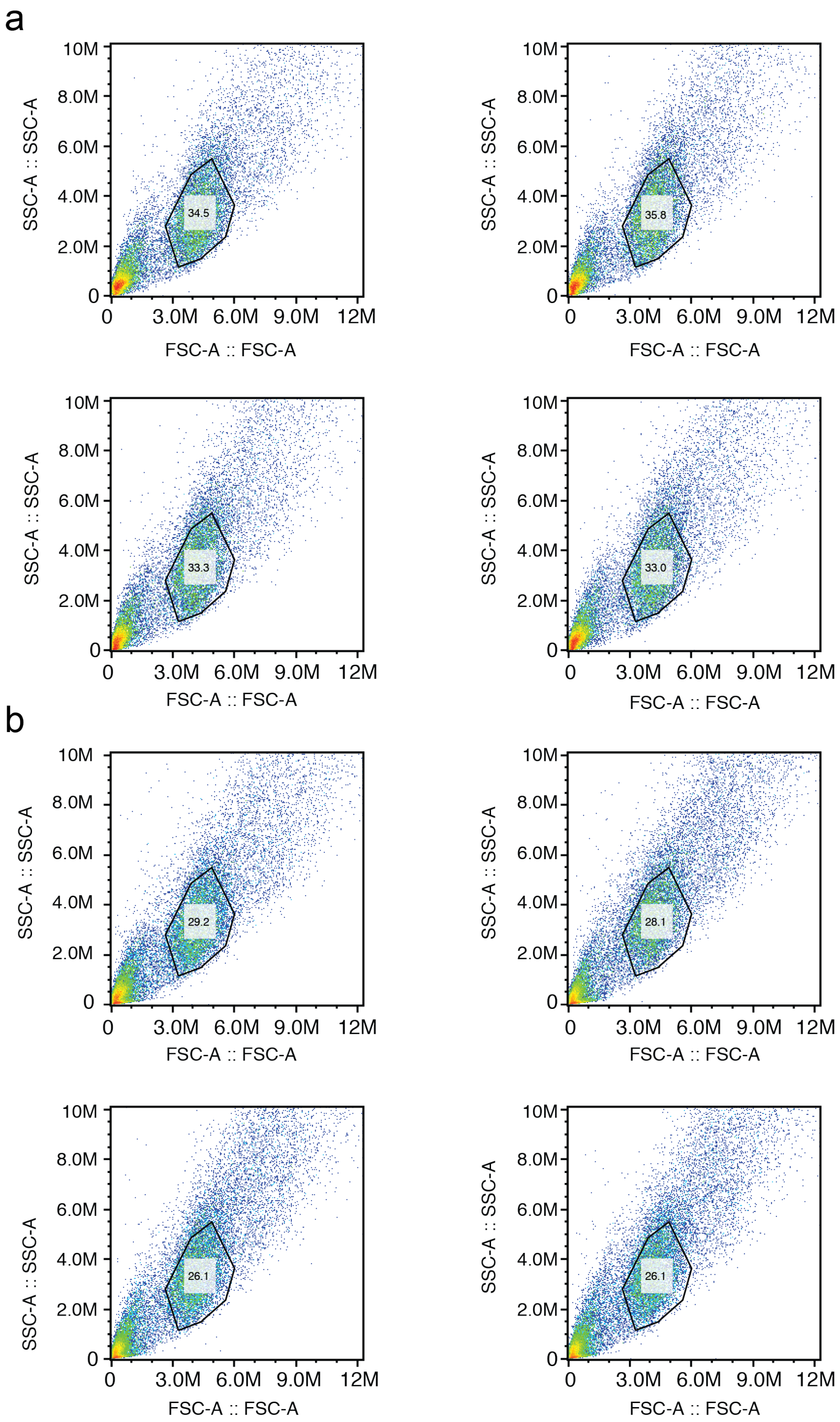

**Supplementary Figure 7.** Flow cytometry analysis of CADH2 levels at the surface of HEK293T cells in the absence (red) or presence (blue) of pervanadate.

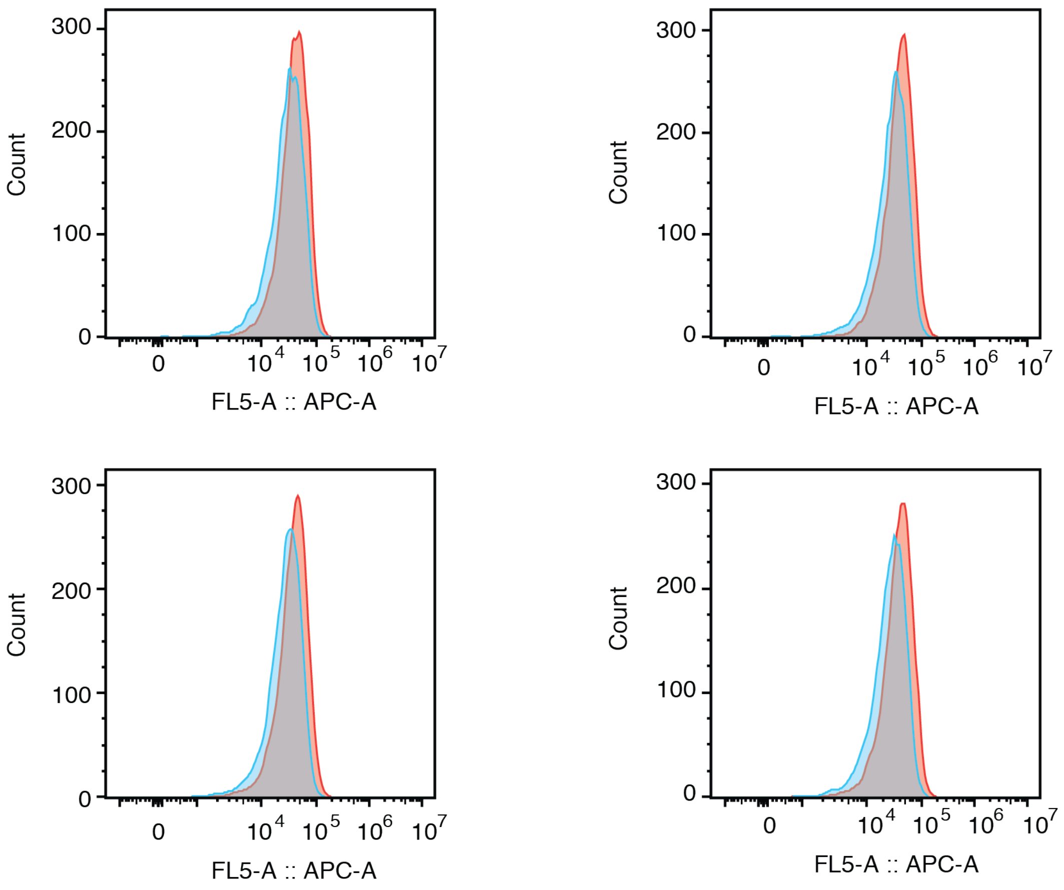

**Supplementary Figure 8.** Flow cytometry gating used for analysis of CD166 levels on the surface of HEK293T cells in the (a) absence or (b) presence of pervanadate.

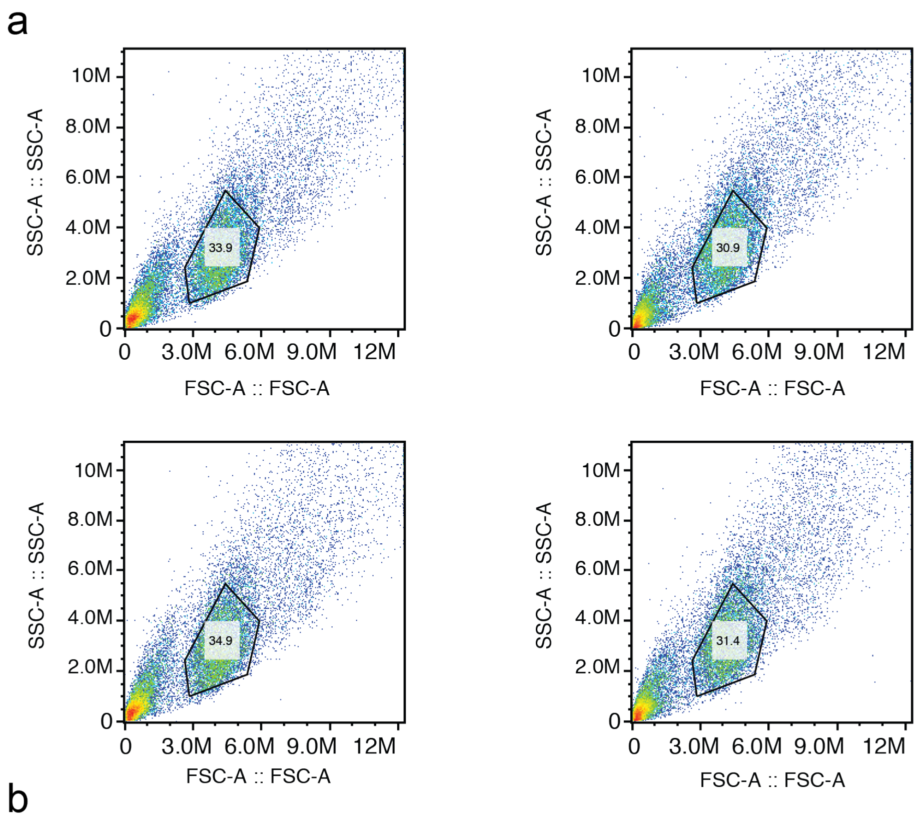


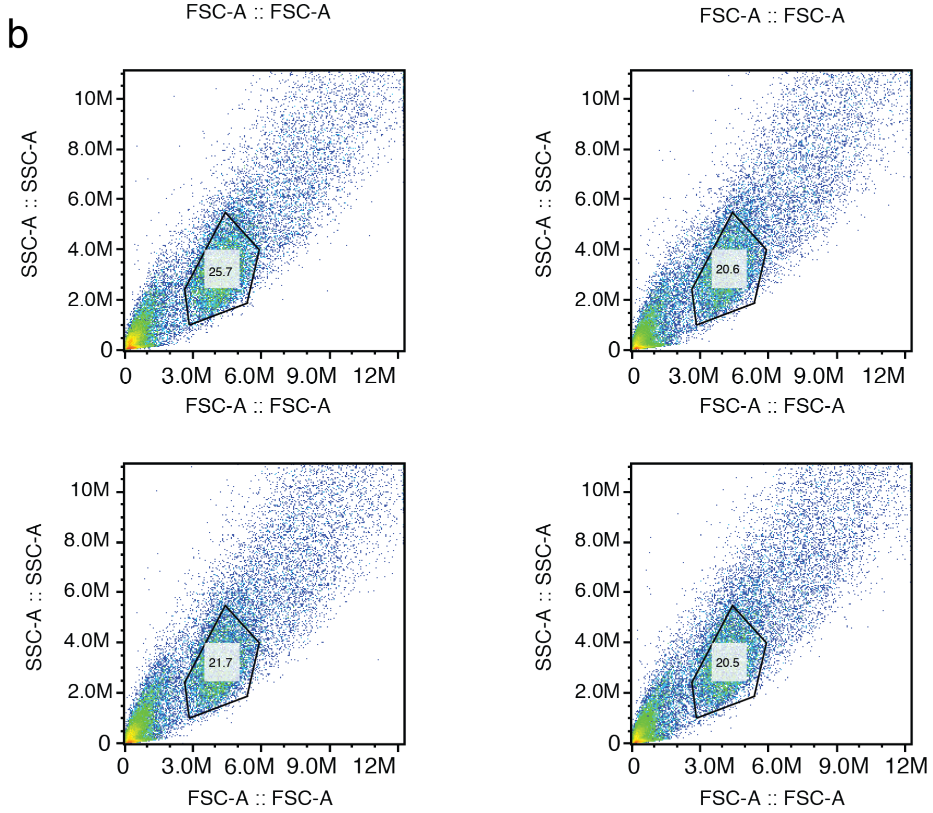

**Supplementary Figure 9.** Flow cytometry analysis of CD166 levels at the surface of HEK293T cells in the absence (red) or presence (blue) of pervanadate.

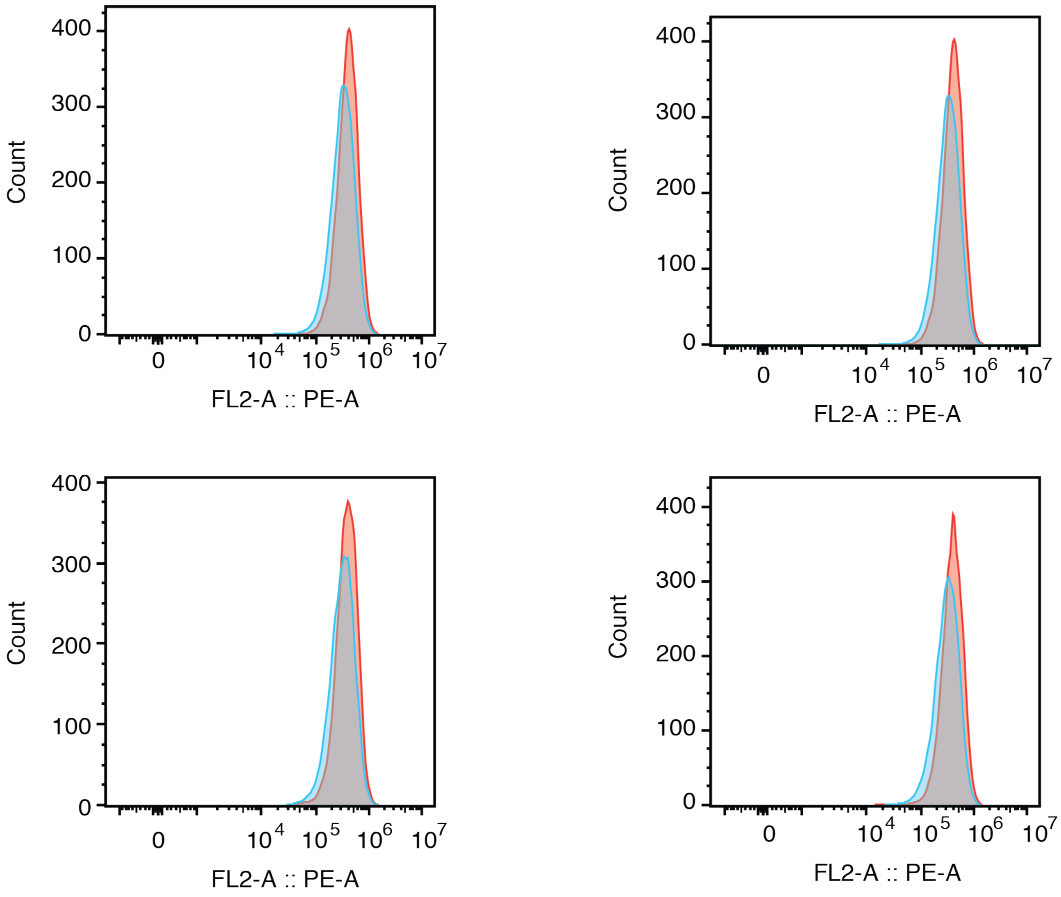

**Supplementary Figure 10.** Flow cytometry gating used for analysis of CD99 levels on the surface of HEK293T cells in the (a) absence or (b) presence of pervanadate.

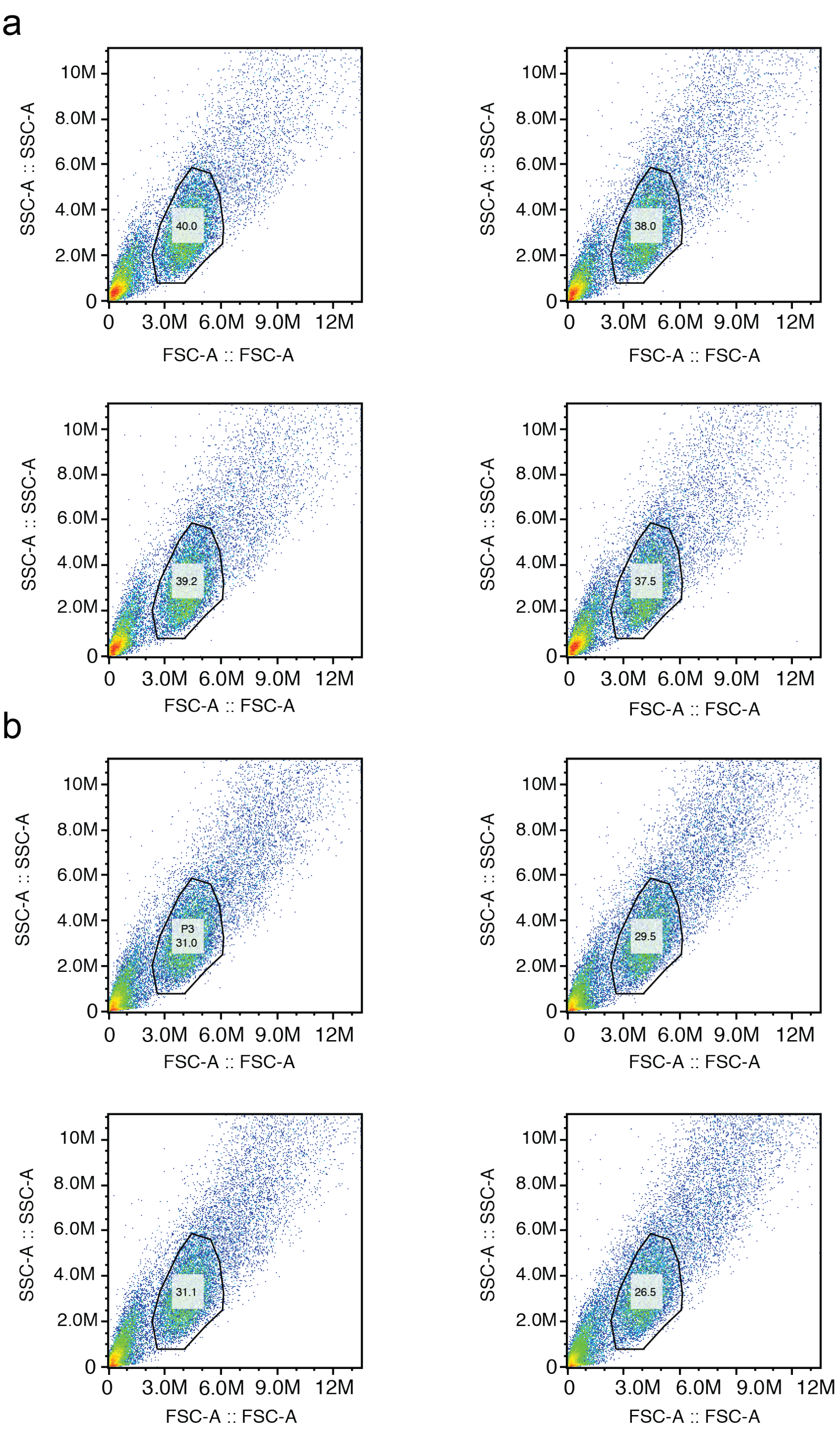

**Supplementary Figure 11.** Flow cytometry analysis of CD99 levels at the surface of HEK293T cells in the absence (red) or presence (blue) of pervanadate.

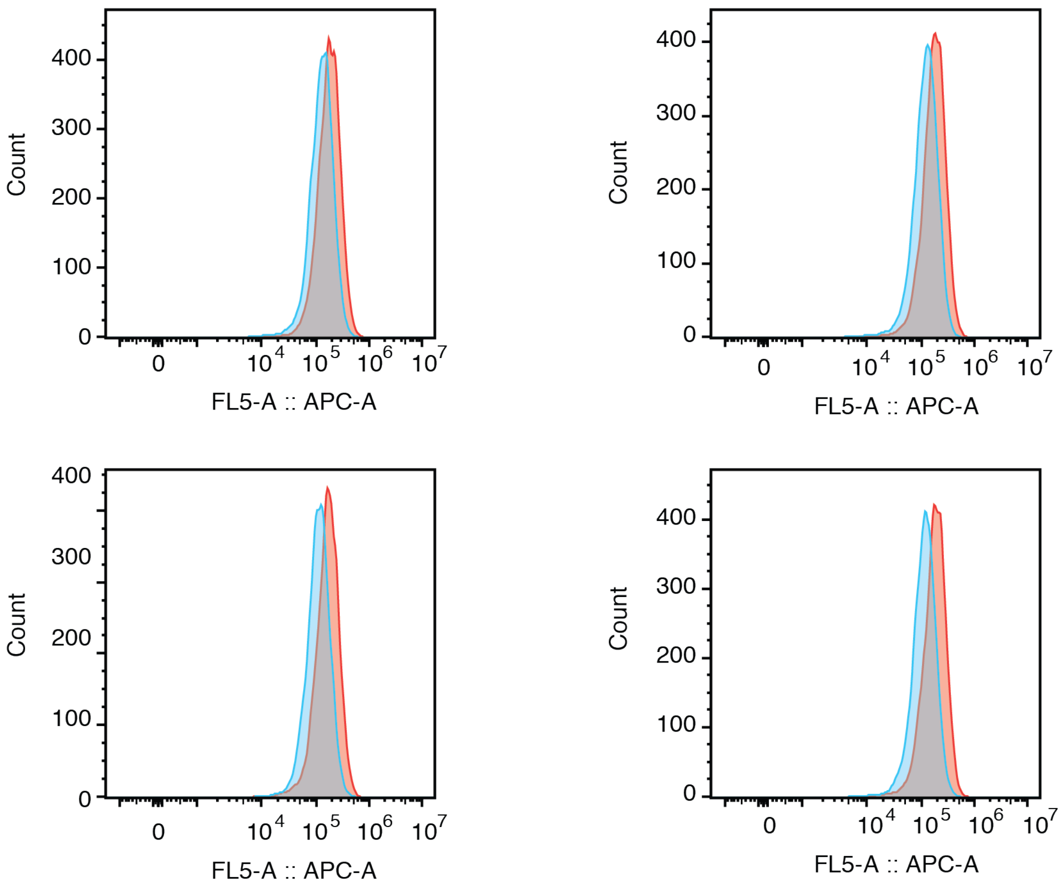
